# PRMT5 as an Epigenetic Target for Group 3 (MYC-driven) Medulloblastoma

**DOI:** 10.64898/2026.04.09.717536

**Authors:** Devendra Kumar, Ajay Sharma, Amit K. Dash, Ranjana Kanchan, Ling Ding, Yashpal S. Chhonker, Sushil Shakyawar, Chittibabu Guda, Ganesh Naik, Daryl J. Murry, Sutapa Ray, Hamid Band, Don W. Coulter, Nagendra K. Chaturvedi

## Abstract

**Background:** Group 3 (MYC-driven) medulloblastoma (MB) is a highly aggressive brain tumor with poor-prognosis and limited treatment options. We previously identified protein-arginine methyltransferase-5 (PRMT5) as a promising target in Group 3 MB with its control on MYC protein stability. In this follow up study, we further mechanistically investigated PRMT5 control on MYC transcription and targeted it pharmacologically for therapeutic proof-of-concept.

**Methods:** Using pharmacogenetic inhibition approaches against PRMT5 in MYC-amplified (Group 3) MB cell line and neurosphere models *in vitro* and *in vivo*, we investigated molecular mechanism(s) and anti-cancer efficacy of PRMT5 inhibition.

**Results:** Our experiments demonstrated that PRMT5 epigenetically regulates MYC transcription in MYC-amplified MB cells by binding to the proximal-promoter region of the MYC gene and contributing to the enriched symmetric-dimethylation of histone H4R3 in the same region. We further showed that PRMT5 is recruited to the MYC promoter by its interaction with BRD4, the major BET-protein responsible for MYC transcription. PRMT5 inhibition caused the suppression of MYC-induced transcriptional programs and target genes, with widespread disruption of splicing across the transcriptome, particularly affecting metabolism-related gene products. Pharmacologic inhibition of PRMT5 using a panel of selective small-molecule inhibitors demonstrates suppression of cell growth/survival in a MYC-dependent manner in MB cells. Moreover, our *in vivo* analyses of PRMT5 inhibition, in mice treated with one of the potent pharmacologic inhibitors, particularly a lipid-decorated form of it, demonstrated reduced cerebellar tumor growth with suppressed MYC expression and prolonged survival of mice with MYC-amplified MB xenografts.

**Conclusions:** Our findings establish a functional link between PRMT5 and MYC-mediated transcriptional regulation, suggesting a promising therapeutic approach targeting the PRMT5-MYC axis for MYC-driven MB.

**Key Points:**

- PRMT5 acts as an epigenetic regulator of MYC transcription, RNA splicing and associated energy metabolism in MYC-driven MB.
- PRMT5 inhibition selectively suppresses cell growth/survival in MYC-driven MB.
- PRMT5 inhibition reduces tumor burden and prolongs survival in a MYC-driven MB mouse model.

**Importance of the Study:** Group 3 medulloblastoma is a highly aggressive pediatric brain tumor marked by MYC amplification, malignant clinical behavior, and poor survival outcomes despite intensive multimodal therapy. Because MYC remains largely undruggable, there is an urgent need for effective and less toxic treatment options for affected children. This study identifies protein arginine methyltransferase 5 (PRMT5) as a key epigenetic regulator of MYC transcription and MYC-dependent oncogenic programs in Group 3 MB. We show that PRMT5 is recruited to the MYC promoter via BRD4, sustains MYC-driven transcription and RNA splicing networks associated with metabolism, and supports MB tumor growth. Importantly, pharmacologic inhibition of PRMT5 using a selective brain-penetrant inhibitor suppresses MYC expression, reduces cerebellar tumor burden, and prolongs survival in MYC-amplified MB models. These findings provide a strong translational rationale for PRMT5 inhibition as a targeted therapeutic strategy for high-risk MB, with the potential to improve outcomes while reducing treatment-related toxicity.

## INTRODUCTION

Medulloblastoma is the most frequent malignant pediatric brain tumor, representing nearly one-fifth of childhood brain cancers. ^1^. Current MB therapies involve surgical resection, radiation and intensive chemotherapy, but fail to cure approximately one-third of patients ^2^. Even among responders, the treatments incur enormous toxicities and are associated with long-lasting side effects. MB has four major molecularly distinct subgroups that include wingless (WNT), Sonic Hedgehog (SHH), Group 3, and Group 4 ^3–5^. Group 3 medulloblastoma frequently demonstrates MYC amplification or overexpression and has the worst clinical outcome among MB subgroups, with survival rates below 60%. MYC-driven medulloblastoma displays aggressive metastatic behavior and is commonly resistant to even intensive multimodal treatment approaches^6–8^. Thus, elucidating mechanisms of MYC-driven MB tumor progression/recurrence and identifying molecular targets suitable for rational targeted therapy are critical to devising more effective and safe regimens for these high-risk patients.

Epigenetic deregulation has emerged as a key driver in MB tumorigenesis, particularly in Group 3 and Group 4 MBs where germline mutations in known cancer predisposition genes are rare ^9,10^. Indeed, dysregulation of epigenetic/chromatin modifiers, including histone methylation/ acetylation marks, is much more common in Group 3 and 4 MBs than in other subgroups ^10–12^. Protein arginine methylation represents a critical post-translational modification that modulates gene transcription and signal transduction. The enzymes responsible for such methylation are the protein arginine methyltransferases (PRMTs). Of these, PRMT5 is the major type II PRMT that regulates oncogenesis-relevant processes such as cell proliferation and differentiation. PRMT5 regulates these cellular processes by modifying gene expression via symmetrical dimethylation of arginine residues in histones H4R3, H3R2, H3R8, and H2AR3 and post-translational regulation of non-histone proteins ^13–15^. The importance of arginine methylation by PRMTs in cancer progression is becoming apparent ^16^. Specifically, PRMT5 is overexpressed in a wide range of cancers, and this often correlates with poor patient prognosis ^17^. PRMT5 overexpression in cancers is thought to cause epigenetic silencing of tumor suppressor and cell cycle genes ^18,19^. Beyond histone substrates, PRMT5 post-translationally methylates key oncogenic transcription factors, including p53, NF-κB (p65), and MYCN ^20–22^. Recent findings implicate PRMT5 in the regulation of aberrant MYC activity in diverse cancers, including glioblastoma^23–25^. Consequently, oncogenic functions have been attributed to PRMT5 and it has recently received attention as a potential therapeutic target. To this end, several selective and potent small-molecule inhibitors have been developed against PRMT5, and their anti-tumor effects are now being assessed in preclinical models and in phase I/II trials ^26,27^.

We previously reported for the first time that PRMT5 is a critical regulator of MYC oncoprotein in MYC-driven Group 3 MB ^28^. We showed that high levels of PRMT5 not only mirror MYC expression but also correlate with poor outcomes in Group 3 MB patients. Mechanistically, we showed that PRMT5 stabilizes MYC protein by physically interacting with it ^28^, raising the intriguing possibility that PRMT5 can regulate MYC function at both the transcription and post-translation levels. However, the exact MYC oncogenic programs regulated by PRMT5 in MB are largely unexplored. In the current study, we explored further mechanistic insights into PRMT5 and MYC interaction as a driver of MB aggressiveness. In addition, our *in vitro* and *in vivo* analyses of the pharmacologic inhibition of PRMT5 demonstrated the potential of the PRMT5-MYC axis as a novel therapeutic target in MB.

## METHODS

### Cell lines and their maintenance

MYC-amplified MB cell lines D-341 and HD-MB03 were obtained from ATCC and DSMZ (Germany), respectively. The SHH-subtype, non-MYC-amplified ONS-76 MB cell line was purchased from Sekisui XenoTech (USA). Group 3 MYC-amplified PDX MB cell lines MED-114FH and MED-411-FH were provided by the Brain Tumor Resource Laboratory at Seattle Children’s Hospital. Normal human astrocyte (NHA) cells were obtained from Lonza.

All cell lines were authenticated by the providing repositories using short tandem repeat profiling and were verified to be mycoplasma-negative using the MycoSensor PCR assay kit (Agilent Technologies). ONS-76, D-341, and HD-MB03 medulloblastoma cell lines were cultured in RPMI-1640 medium supplemented with 10% heat-inactivated fetal bovine serum and 1% penicillin–streptomycin in a humidified incubator at 37 °C with 5% CO₂. PDX MB cell lines MED-114FH and MED-411FH were grown as spheroids in non-coated plates using NeuroCult NS-A medium supplemented with proliferation supplements (Stemcell Technologies) and 1% penicillin–streptomycin in a humidified incubator at 37 °C with 5% CO₂. Only cells at fewer than 10 passages were used for all experiments.

### PRMT5 inhibitors

Four PRMT5 inhibitors (EPZ015666, GSK332595, LLY-283, JNJ64619178) were purchased from MedChemExpress LLC or Selleckchem Company. Stock solutions of inhibitors (1 mM or 10 mM) were prepared in dimethyl sulfoxide (DMSO) and stored at −20 °C. For all experiments, the DMSO concentration was matched to that of the highest drug dose and used as a solvent control.

### Cell growth, apoptosis, and cell cycle analyses

The effects of inhibitor treatment on MB cell growth, apoptosis, and cell cycle progression were analyzed using MTT assays, Annexin V staining, and propidium iodide staining, respectively, following established protocols ^29–31^.

### Western blotting

Western blot analyses were performed using protocols previously established in our laboratory ^29^. Primary antibodies used in this study included MYC (Cell Signaling Technology #18583), PRMT5 (Cell Signaling Technology #79998), SDMA (Cell Signaling Technology #13222), H4R3me2s (Abcam, ab5823), H3R8me2s (Abcam, ab130740) and GAPDH (Cell Signaling Technology #5174).

### Quantitative RT-PCR (qRT-PCR)

RNeasy Kit (Qiagen) was used for isolating total RNA, and 2 µg of RNA was reverse-transcribed into cDNA using the SuperScript Verso cDNA synthesis kit (Promega). Quantitative real-time PCR was conducted in 10 µL reactions using SYBR Green Supermix and gene-specific primers for MYC, PRMT5, and GAPDH (Applied Biosystems). Reactions were performed using QuantStudio 3 Real-Time PCR System, and data were analyzed with QuantStudio software (Applied Biosystems).

### siRNAs and transfection

Scrambled control (sc-37007), PRMT5 (sc-41073), and BRD4 (sc-43639) siRNAs were obtained from Santa Cruz Biotechnology (Dallas, TX, USA). siRNAs were resuspended in RNase-free water to a stock concentration of 10 µM and stored at −20 °C. MB cells were transiently transfected with either control or pooled gene-specific siRNAs (three 19–25 nt siRNAs; 50 nM final concentration) using Lipofectamine 2000 (Invitrogen) following the manufacturer’s instructions. Seventy-two hours post-transfection, cells were collected for downstream analyses.

### Luciferase-based transcriptional activity assay

Overall transcriptional activity of MYC in PRMT5 knock-down HD-MB03 cells was assessed by transiently transfecting MYC promoter tagged with luciferase (luc) reporter gene (Creative Biomart #1785) using a bioluminescence-based dual-luciferase reporter assay kit (Promega # E1910) according to instructions.

### Co-immunoprecipitation

For co-immunoprecipitation experiments, 500 µg of HD-MB03 cell lysate was precleared with protein A–Sepharose beads (Cell Signaling Technology) for 1 h at 4 °C and incubated with 8 µg of PRMT5 antibody overnight at 4 °C. Immune complexes were collected by incubation with protein A–Sepharose beads for 2 h at 4 °C with rotation. Following PBS washes, beads were resuspended in 1× Laemmli buffer and subjected to western blot analysis.

### Chromatin-immunoprecipitation assay

Using a previously described protocol, ChIP assay was performed using PRMT5, H4R3me2s, and BRD4 antibodies ^32^. Genomic qPCR was performed in a Quant Studio 3 system, using SYBR green. ChIP primers for human MYC promoters (14905) were purchased from Cell Signaling Technology. The sequences of MYC primers used were forward primer (5′->3′): TGAGTATAAAAGCCGGTTTTCG; reverse primer (5′->3′): CTGCCTCTCGCTGGAATTACTA. Data are normalized to either percent input DNA or IgG control.

### Sphere assay

MED-114FH and MED-411FH MB cells (20,000 cells per well) were resuspended in NeuroCult medium and seeded into non-coated 6-well plates to allow spheroid formation for 72 h. Formed spheroids were treated with PRMT5 inhibitors for an additional 72 h, after which aggregates larger than 50 µm were quantified and imaged using an EVOS Auto Imaging System (Life Technologies). Spheroids were then collected for western blot analysis of MYC protein levels.

### Metabolome Analysis

HD-MB03 cells were cultured in RPMI medium containing vehicle or the PRMT5 inhibitor JNJ64619178 (1 µM) for 24 h. Cells were subsequently rinsed and subjected to extraction with 80% ice-cold methanol on dry ice for polar metabolite collection. Targeted metabolomic data analysis was performed as previously reported. ^33^.

### Seahorse analyses

The oxygen consumption rate (OCR) and extracellular acidification rate (ECAR) of the cells were assessed using the Seahorse XFe24 Analyzer (Agilent) according to the manufacturer’s instructions. HD-MB03 cells (4 × 10³ per well) were seeded in Seahorse XFe24 24-well assay plates and incubated overnight at 37 °C to allow attachment. Cells were subsequently treated with vehicle control or 1 µM JNJ64619178 for 24 h. Prior to measurement, culture medium was replaced with Seahorse assay medium supplemented with 10 mM glucose, 1 mM sodium pyruvate, and 2 mM glutamine (pH 7.4), and cells were incubated at 37 °C for 1 h. Basal ECAR and OCR were recorded for 24 min, followed by glycolytic and mitochondrial stress tests. Data were plotted on a metabolic grid to depict degrees of aerobic and glycolytic activities.

### RNA sequencing and gene expression analyses

RNA from PRMT5 inhibitor-treated HD-MB03 cells was isolated using the Qiagen RNeasy Kit. RNA integrity and sequencing quality were verified with an Agilent 2100 Bioanalyzer. High-quality RNA was used to generate sequencing libraries with the TruSeq RNA Sample Prep V2 Kit, followed by next-generation sequencing on the Illumina NextSeq550 platform at the UNMC Genomics Core Facility. All samples were processed in triplicate. Raw FASTQ files were analyzed using a standardized pipeline in which STAR was used for alignment and RSEM for gene and isoform-level annotation and quantification. Normalized FPKM and TPM values were calculated for all detected genes, and differential expression and gene-set enrichment analyses (GSEA) between treatment groups were performed by the UNMC Bioinformatics Core Facility. To evaluate global transcriptional alterations, we conducted Gene Ontology (GO) enrichment and Gene Set Enrichment Analysis (GSEA). GO Biological Process enrichment was performed using standard over-representation analysis to identify functional categories significantly enriched among differentially expressed genes meeting the criteria of log2(fold-change) ≥ 1 and p < 0.05. GSEA was performed using a pre-ranked list of all expressed genes ordered by fold-change, enabling the identification of coordinated pathway-level alterations without applying an arbitrary significance threshold. Enrichment scores and normalized enrichment scores (NES) were generated using permutation testing, and pathways with a false discovery rate (FDR) < 0.25 were considered significantly enriched.

To analyze RNA splicing, we first assessed RNA-seq read quality using FastQC and summarized the reports with MultiQC. Reads passing quality control were aligned to the reference genome with STAR, and gene-level counts were generated using featureCounts. Differential gene expression analysis was then performed in R with edgeR, applying TMM normalization and filtering out genes expressed at low levels before statistical testing. To specifically examine alternative splicing, transcript abundances were estimated with Kallisto, followed by SUPPA2-based annotation of splicing events. Changes in percent spliced-in (ΔPSI) values and their statistical significance were calculated across conditions. The analysis focused on major splicing event types, including skipped exons (SE), alternative 5′ and 3′ splice sites (A5/A3), mutually exclusive exons (MX), retained introns (RI), and alternative first and last exons (AF/AL).

### LNP formulation and brain tissue distribution

Lipid nanoparticles (LNPs) are efficient vehicles for hydrophobic drugs due to their excellent biocompatibility, physical stability, lipid-based internal structure, and easy surface modification. Emulsion is one of the early LNP models. Lipid nanoparticles (LNPs) of JNJ64619178 were prepared using biodegradable and biocompatible self-emulsifying drug delivery system (SEDDS) formulations, comprised of surfactants (Labrafil M 2125 CS and Kolliphor RH40), oil phase (Capryol 90) and PEG400, as described previously ^34^. JNJ64619178 loaded SEDDS was prepared by dissolving JNJ powder into the blank formulation, followed by dilution in PBS to form an oil-in-water emulsion.

Brain tissue distribution of free or formulated JNJ64619178 (10 mg/kg) was evaluated in balb/c mice following oral administration. Animals were divided into three groups and euthanized at 2, 8, and 24 hours post dose. To assess brain penetration, transcranial perfusion with phosphate-buffered saline was performed prior to brain collection. The concentration of free or formulated JNJ64618178 in the brain was analyzed using LC-MS.

### Animal studies

All animal studies were performed in accordance with protocols approved by the University of Nebraska Medical Center (UNMC) Institutional Animal Care and Use Committee (IACUC). For subcutaneous xenograft experiments, six- to eight-week-old female NSG mice (The Jackson Laboratory) were injected subcutaneously in the right flank with 2.5 × 10⁵ HD-MB03 medulloblastoma cells resuspended in 100 µL of PBS and combined with Matrigel at a 1:3 ratio. Once tumors became palpable, approximately 10 days after implantation, mice were randomized into two treatment groups (vehicle or JNJ64619178; n = 5 per group) and treated for three weeks. The vehicle formulation consisted of 5% DMSO, 45% PEG 300, and 2% Tween 80, while JNJ64619178 was administered orally at 10 mg/kg, five times per week. Tumor growth was monitored twice weekly using digital caliper measurements. Mice were euthanized by CO₂ inhalation when tumor volumes approached 2 cm³, and tumors along with major organs were collected for histological and immunohistochemical evaluation. Immunohistochemistry was performed following a previously established protocol in our laboratory using rabbit antibodies against MYC (1:500) and Ki-67 (1:500) obtained from Abcam.

For orthotopic xenograft experiments, 1 × 10⁵ HD-MB03 cells suspended in 5 µL of PBS were stereotactically injected into the cerebella of NSG mice using a digital stereotaxic system (Stoelting Instruments). Ten days after implantation, mice were randomly assigned to three treatment groups—vehicle, JNJ64619178, or LNP-formulated JNJ64619178 (n = 5 per group)—and treated according to the regimen used for subcutaneous xenograft studies. Mouse survival was monitored daily, with severe ataxia, tumor-associated morbidity, or body weight loss exceeding 20% defined as humane endpoints. Cerebellar tumors were subsequently analyzed by histological (H&E) staining and MYC immunohistochemistry.

## RESULTS

### MYC is an epigenetic target of PRMT5 in MB cells

We showed previously that PRMT5 forms a complex with MYC, leading to MYC protein stabilization in MYC-amplified MB cells ^28^. However, it is unexplored if PRMT5 also regulates MYC transcription, which is particularly relevant because MYC-induced transcriptional programs play key roles in tumorigenesis. To further determine how PRMT5 regulates MYC transcription, we examined the effect of PRMT5 inhibition on MYC transcription by quantitative real-time PCR (qRT-PCR) and observed that transient knockdown of PRMT5 in MYC-amplified MB cells decreased the mRNA level of MYC by ∼50% **(Figure 1A)**. As PRMT5 may regulate MYC transcription epigenetically, we examined the effect of PRMT5 knockdown on the MYC-Luciferase reporter gene (MYC-Luc) activity and observed that PRMT5 knockdown had a significant inhibitory effect on the MYC-Luc activity **(Figure 1B)**. Thus, it likely acts through epigenetic control of MYC transcription. Indeed, PRMT5 and its symmetric dimethylation activity on H4R3 (H4R3me2s) were significantly enriched on the proximal promoter region of the MYC gene **(Figure 1C)**. Consistently, the knockdown of PRMT5 led to a significantly decreased enrichment of both PRMT5 and H4R3me2s to the MYC promoter region **(Figure 1D and E)**. Together, these results demonstrate that PRMT5 epigenetically activates MYC transcription by symmetrically dimethylating H4R3 in MYC-amplified MB cells.

**Figure 1.**
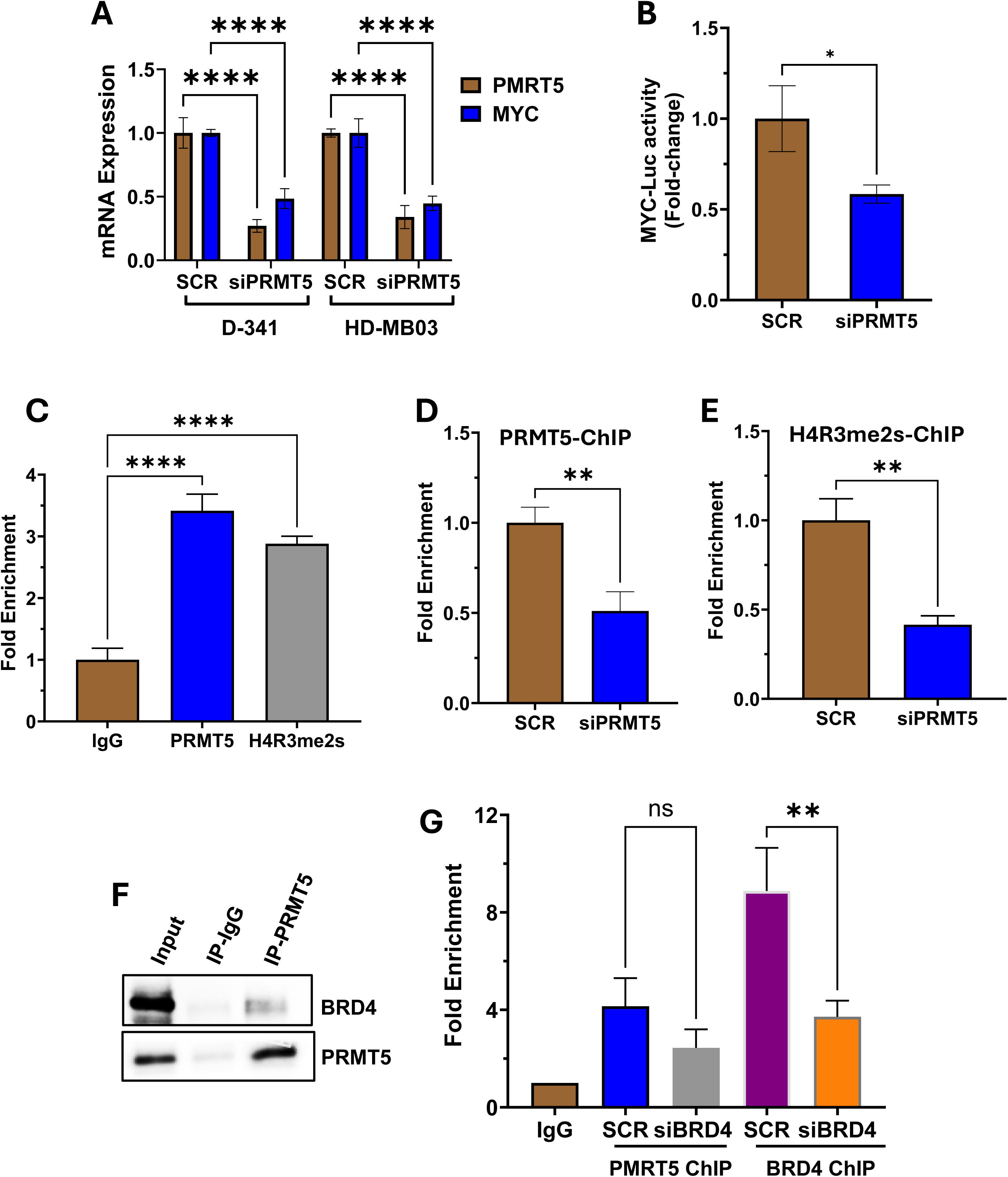
Epigenetic activation of MYC transcription by PRMT5 in MYC-amplified MB cells. **(A)** qPCR analysis for the expression of PRMT5/MYC mRNA in two MYC-amplified cell lines with transiently knocked-down of PRMT5 (using siRNAs) at 72 h. ****, p<0.0001 (student t test, SCR vs siPRMT5). **(B)** Effect of PRMT5 knockdown (siRNAs) on MYC-luciferase reporter gene (MYC-Luc) activity in HD-MB03 cells. *, p<0.05 (student t test) **(C)** ChIP analyses for the enrichment of PRMT5 and H4R3me2s on the proximal promoter region of the MYC gene in HD-MB03 cells. ****, p<0.0001 (student t test). ChIP analyses for the enrichment/binding of PRMT5 **(D)** and H4R3me2s **(E)** to the proximal promoter region of MYC gene, in PRMT5 knockdown (siRNAs) HD-MB03 cells. (**F**) Co-immunoprecipitation of BRD4 with PRMT5. (**G**) ChIP analyses for the enrichment/binding of PRMT5 and BRD4 to the proximal promoter region of MYC gene, in BRD4 knockdown (siRNAs) HD-MB03 cells. *, p<0.05; **, p<0.05 (student t test).

To determine how PRMT5 is recruited to the MYC promoter, we examined whether PRMT5 interacts with bromodomain-containing protein 4 (BRD4), the well-characterized epigenetic regulator that binds to the MYC gene promoter and positively regulates MYC transcription ^35^. Indeed, BRD4 was co-immunoprecipitated with PRMT5 in HD-MB03 cells **(Figure 1F)**. Next, we confirmed that the knockdown of BRD4 indeed repressed MYC expression. Also, the knockdown of BRD4 in this cell line not only abolished the binding of BRD4 to the proximal promoter region of the MYC gene but also abolished the binding of PRMT5 to the same region **(Figure 1G)**. These results together suggest that BRD4 and PRMT5 form a complex on the MYC proximal promoter region to activate MYC transcription.

### PRMT5 inhibition suppresses MYC-induced transcriptional programs and alternate splicing in MB cells

To explore the mechanism(s) of PRMT5 targeting in an unbiased manner, we performed a transcriptomic analysis (RNA-seq) in the well-characterized HD-MB03 cells treated with JNJ64619178 (1 µM) for 24 h. Using log2(fold-change) ≥1 with p-value < 0.05, we detected 2015 upregulated and 2027 downregulated genes (of all 12,985 detectable genes) by JNJ64619178 **(Figure 2A)**, suggesting that targeting PRMT5 may effectively modulate genome-wide transcription in MYC-driven MB cells. An unbiased screen of pathways altered by JNJ64619178 (using GO-BP classification and gene-set enrichment analysis) identified significant alteration of 20 pathways **(Figure 2B)**. Of these, a significant number of tumorigenic pathways, particularly MYC-target pathways, were down-regulated **(Figure 2C)**. These altered pathways included MYC-target signatures, mRNA splicing, several metabolic pathways, chromatin modification, cell cycle and DNA repair **(Figure 2B and C)**. PRMT5 is critical for spliceosome assembly, and tumor cells are quite dependent on a hyperactive core splicing machinery ^36^. To broadly assess whether splicing disruption occurred on JNJ64619178 treatment, we analyzed global alternative splicing (AS) changes in our RNA-seq dataset. We identified 1280 AS events in protein-coding genes, with the majority being skipped exons and alternative first exons **(Figure 2D)**, two events that have shown PRMT5 dependency in other tumor types ^37,38^. It is expected that each ASE should be equally likely to be included or excluded after treatment. GO analysis of the altered splicing events revealed significant enrichment in genes involved in RNA splicing and several categories related to metabolic pathways **(Figure 2E)**. Among the aberrantly spliced genes, we identified genes, *GGT6* and *GPT2*, that are critically involved in glutamate metabolism ^39^ and *C1QBP*, a mitochondrial protein essential for oxidative phosphorylation and inflammation ^40^ **(Figure 2F)**.

**Figure 2.**
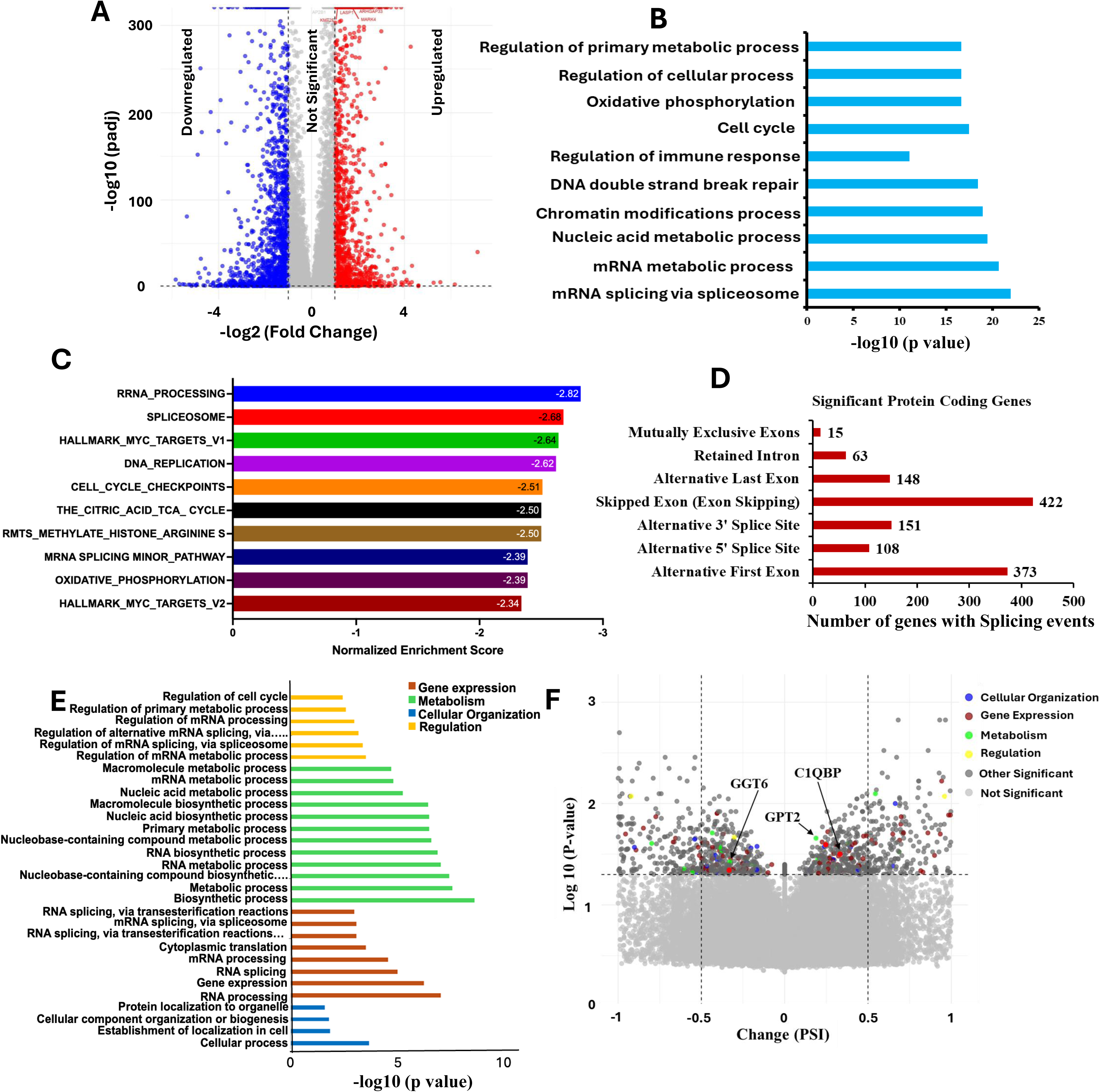
Effects of JNJ64619178 (JNJ) on global gene expression and splicing. RNA-sequencing was used to assess global gene expression changes in HD-MB03 cells 24 h after treatment with DMSO (vehicle) or JNJ64619178 (1 µM). **(A)** Volcano plot displaying genes significantly upregulated or downregulated in response to PRMT5 inhibition. **(B)** Gene Ontology Biological Process (GO-BP) functional analysis for top 10 pathways altered by JNJ. **(C)** Gene sets enrichment analysis (GSEA) (with p<0.01 and FDR<0.2) for top 10 pathways/gene sets (including MYC-associated target gene sets, RNA splicing, metabolism) altered by PRMT5 inhibition. **(D)** Number of the aberrant splicing events in protein coding genes disrupted by PRMT5 inhibition. **(E)** GO-BP functional analysis for the aberrant splicing events altered by PRMT5 inhibition. **(F)** Volcano plot represents all the significant (p<0.05) splicing events. Selected genes from the indicated GO-BP functional classification are highlighted. Metabolism associated genes *GGT6, GPT2* and *C1QBP1* are identified.

### PRMT5 inhibition alters energy metabolism in MYC-driven MB cells

Given such a prominent role of PRMT5 in regulating gene expression and splicing associated with cancer metabolism, we next wanted to see whether PRMT5 inhibition modulates overall metabolism in MYC-amplified MB cells. Targeted metabolomic profiling was performed using LC–MS/MS in control and PRMT5 inhibitor–treated (JNJ64619178) HD-MB03 cells. A panel of 173 standard metabolites representing major cellular metabolic pathways was analyzed. Metabolomic and pathway analyses were conducted using MetaboAnalyst 5.0 software. Principal component analysis of metabolite profiles from control and JNJ64619178-treated samples, each examined in three technical replicates, revealed clear treatment-dependent metabolic reprogramming (**Supplementary Figure 1**). The PCA plot shows tight clustering of technical replicates, indicating high reproducibility within each condition.

Statistical comparison of metabolite levels between conditions using a student’s t-test (p < 0.05) identified 60 significantly altered metabolites. A heatmap was generated to illustrate the relative abundance patterns of these metabolites across control (DMSO) and JNJ64619178- or LLY-283-treated groups **(Figure 3A; Supplementary Figure 2A)**. Sixty primary metabolites were significantly altered, suggesting pronounced metabolic reprogramming between control and JNJ64619178- or LLY-283-treated groups. Pathway enrichment analysis further demonstrated that each treatment significantly modulated 25 metabolic pathways (**Figure 3B; Supplementary Figure 2B)**. JNJ64619178 treatment resulted in extensive alterations in key energy-related metabolic pathways, including amino acid metabolism (arginine and proline, aspartate, glutamate, and glutathione), pyruvate metabolism, the tricarboxylic acid (TCA) cycle, and mitochondrial electron transport chain activity (**Figure 3B; Supplementary Figure 2B**). Comparable metabolic changes were observed following treatment with LLY-283, another potent PRMT5 inhibitor (**Supplementary Figure 2B)**. Importantly, these affected pathways are well known to be upregulated by MYC activation, supporting the conclusion that PRMT5 inhibition effectively targets MYC-driven metabolic programs.

**Figure 3.**
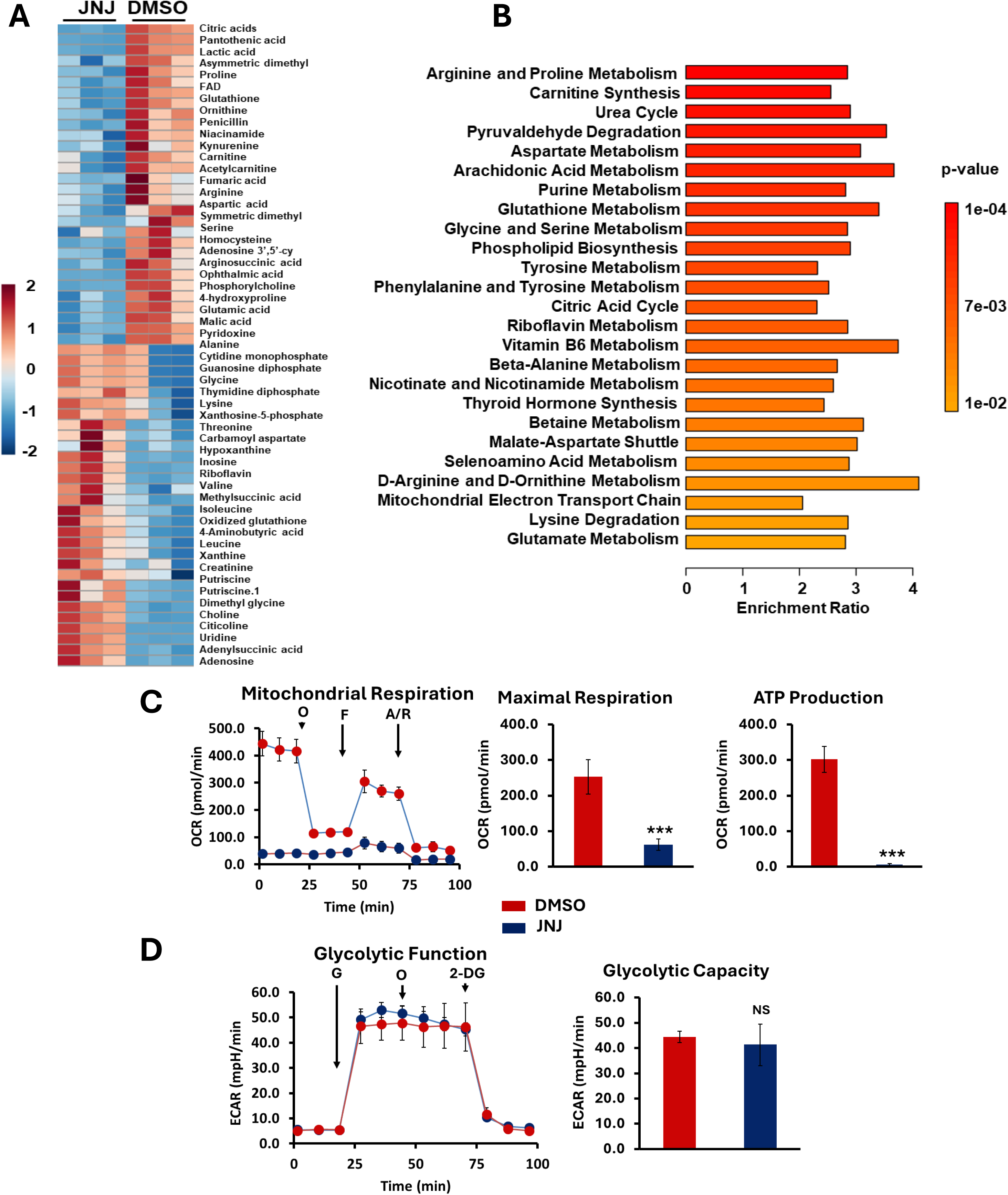
JNJ64619178 (JNJ) treatment alters energy metabolism in MB. **(A)** Heatmap showing top 60 metabolites altered in HD-MB03 cells treated with JNJ (1 µM) in triplicate for 24 h. Color intensity represents the magnitude of alteration in individual metabolites. Scale for color intensity is shown adjacent to the heatmap. **(B)** Pathway analysis showing significantly altered metabolic pathways in JNJ treated HD-MB03 cells, compared to DMSO solvent. Scale under the pathway plot shows the fold enrichment and color scale adjacent to pathway plot indicates significance (*p*-value) for altered pathways. **(C)** Oxygen consumption rate (OCR) analysis for mitochondrial oxidative phosphorylation status in HD-MB03 cells after treatment with 1 µM JNJ for 24 h. O, oligomycin; F, FCCP; A/R, antimycin/rotenone. The bar graphs show maximal respiration and ATP production activities derived from OCR activities shown in line graph. The results represent the mean ± SEM of three replicates. \*\*\**p* < 0.001 (Student t test, DMSO vs JNJ). **(D)** Extracellular acidification rate (ECAR) analysis for glycolytic activities in HD-MB03 cells after treatment with 1 µM JNJ for 24 h. G, glucose; O, oligomycin; 2-DG, 2-deoxyglucose. The bar graph shows glycolytic capacity derived from ECAR activities shown in line graph. The results represent the mean ± SEM of three replicates. NS (not significant).

Targeted metabolic analyses were performed using Seahorse-based assays that evaluated the effects of the PRMT5 inhibitor JNJ64619178 on oxidative phosphorylation and glycolysis in HD-MB03 cells. Oxidative phosphorylation and glycolytic activity were measured in real time using oxygen consumption rates (OCR) and extracellular acidification rates (ECAR), respectively. Treatment with JNJ64619178 led to a significant reduction in mitochondrial maximal respiration, which was accompanied by marked decreases in maximal respiratory capacity and OXPHOS-dependent ATP production **(Figure 3C)**.

In ECAR analysis, no significant effect of JNJ64619178 on glycolysis (glycolytic capacity) was observed, **(Figure 3D),** a result consistent with our metabolome pathway analysis. These results were consistent with another MYC-amplified MB cell line D-425 **(Supplementary Figure 3)**. Together, our results suggest that JNJ64619178 inhibits oxidative phosphorylation activities associated with energy metabolism, but not glycolysis, in MYC-driven MB cells.

### Effect of pharmacological inhibition on MB cell growth, cell cycle and apoptosis in vitro

We tested the therapeutic potential of targeting PRMT5 using a panel of selective small molecule inhibitors (EPZ015666, GSK332595, LLY-283, JNJ64619178) against two MYC-amplified (Group 3) MB cell lines, one non-MYC (SHH) MB cell line, and a normal human astrocyte (NHA) cell line. To examine the anti-MB efficacy of such inhibitors, a cell growth assay (MTT) was performed. MTT results shown in **Figure 4A** clearly demonstrated a dose-dependent cell growth inhibition in MB cell lines by PRMT5 inhibitors. IC50 values demonstrated strong efficacy (low µM potency) of each PRMT5 inhibitor, with significantly lower IC50 values in MYC-amplified MB cell lines **(Figure 4B)**, suggesting PRMT5’s on-target specificity to Group 3 (MYC-driven) MB. PRMT5 inhibitors were not effective in reducing cell growth in normal human astrocyte (NHA) cells, even with high micromolar doses, suggesting no off-target effect of PRMT5 inhibition. Among these inhibitors, JNJ64619178 and EPZ015666 were most potent in inhibiting cell growth/survival of MYC-driven MB cells, with low-µM potency **(Figure 4A and B)**, and this was associated with apoptosis induction and G1-S cell cycle arrest as evident by Annexin-V and propidium-iodide staining assays, respectively **(Figure 4C and D)**. Also, EPZ015666 and JNJ64619178 significantly downregulated the expression of PRMT5 (and its symmetric-dimethylation activity) and MYC protein in MYC-amplified MB (HD-MB03, D-341) cell lines **(Figure 4E)**. Together, results suggest that pharmacologic inhibition of PRMT5 can be a valid approach against MYC-driven MB.

**Figure 4.**
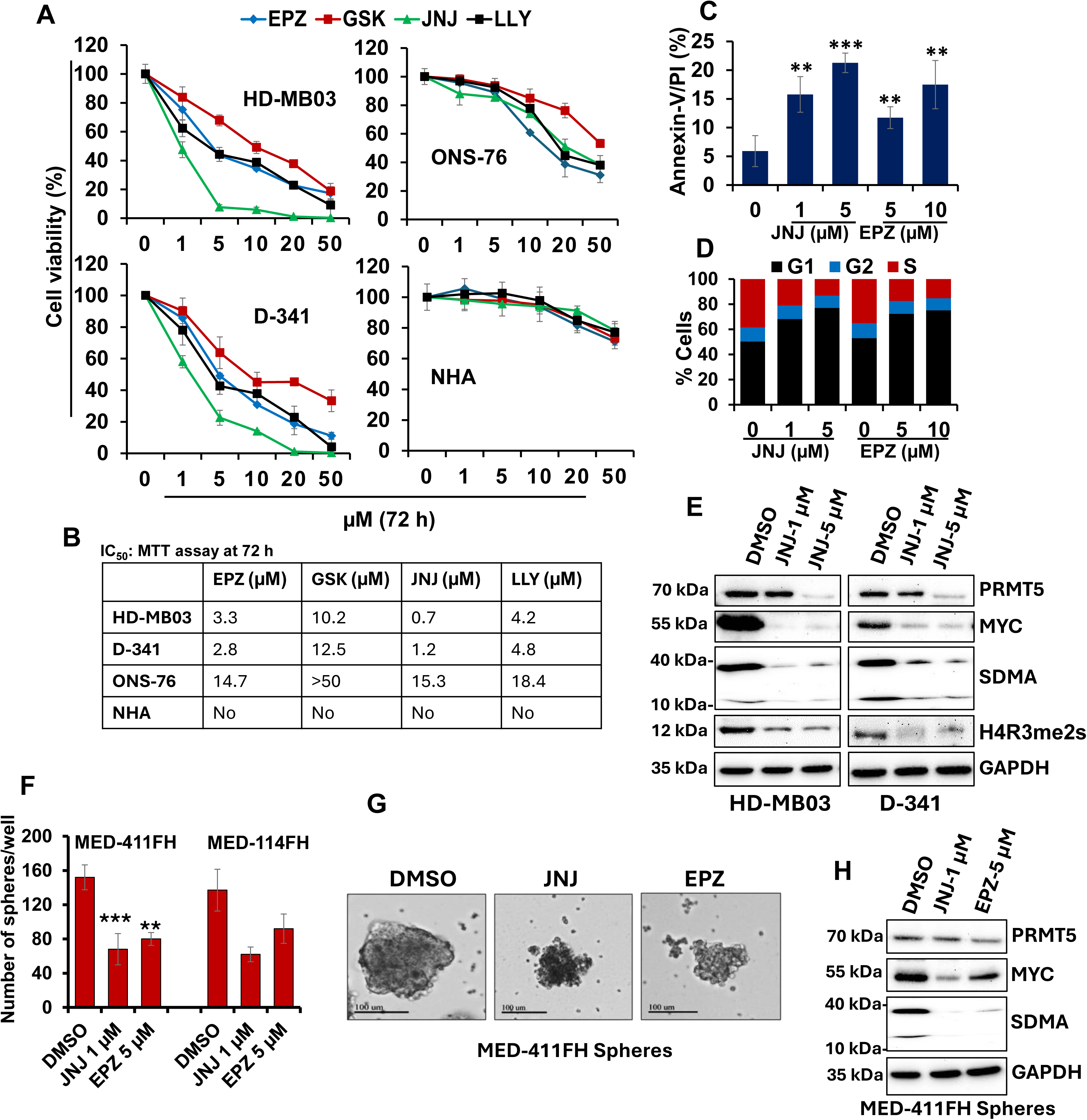
*In vitro* efficacy of PRMT5 inhibitors in MB cell lines. **(A)** MTT assay showing the effects of PRMT5 inhibitors (1-50 µM) on cell growth of MYC-amplified MB (HD-MB03, D341), non-MYC MB (ONS-76) and normal human astrocyte (NHA) cell lines. **(B)** IC_50_ values of PRMT5 inhibitors in the indicated cell lines. **(C)** Annexin-V assay showing effects of PRMT5 inhibitors (JNJ, EPZ) on apoptosis in HD-MB03 cells. **, p<0.01, ***, p<0.001 (relative to ‘0’ (vehicle control)). **(D)** Cell cycle profile in HD-MB03 cells treated with PRMT5 inhibitors (EPZ, JNJ). **(E)** Western blot analysis for the expression of the indicated key proteins in JNJ-treated HD-MB03 cells. (**F)** Quantification of spheres following treatment of JNJ and EPZ in two PDX-derived MB cell lines (MED-411FH, MED-114FH) at 72 h. Values, mean ± SEM. \**p* < 0.05; \*\**p* < 0.01; \*\*\**p* < 0.005; \*\*\**p* < 0.001 (Student-*t*-test). **(G)** Representative sphere (MED-411FH) images showing disruption of spheres in each treatment. **(H)** Western blot results showing the expression of indicated proteins in MED-411 spheres treated with JNJ and EPZ.

We next confirmed the efficacy of PRMT5 inhibition in two previously characterized MYC-amplified Group 3 MB PDX cell lines, MED-114FH and MED-411FH. ^41^. Because these PDX models grow as spheroids, we tested the efficacy of the PRMT5 inhibitors JNJ64619178 and EPZ015666 in spheroid cultures. Established medulloblastoma spheroids cultured in NeuroCult medium were treated with the indicated inhibitors followed by the assessment of sphere numbers and MYC/PRMT5 expression. Both inhibitors showed strong efficacy in the inhibition of sphere counts/formation, the expression of MYC and PRMT5, and the global symmetric dimethylation of arginine (SDMA) in PDX cells **(Figure 4F-H)**, all consistent with anti-cancer efficacy of PRMT5 inhibitors on cultured MYC-amplified Group 3 MB.

### Pharmacologic inhibition of PRMT5 reduces tumor growth in MYC-driven MB xenografts

To evaluate therapeutic potential of PRMT5 inhibition against MYC-driven MB in vivo, we used JNJ64619178 as a potent and clinically relevant inhibitor of PRMT5 ^42,43^. We reported previously that although JNJ64819178 crosses the blood-brain barrier (BBB), only sub-therapeutic brain concentrations were achieved ^44^. Therefore, we formulated this inhibitor with lipid nanoparticles (LNPs) to improve brain penetration. We administered JNJ64619178 (10 mg/kg) with and without LNP formulation to mice through oral gavage, then collected brain samples after 2h, 8h, and 24 h. Our results showed significantly greater brain accumulation of LNP-loaded JNJ64619178 compared to free JNJ64619178 at 2 h (176 ng/g versus 105 ng/g), 8 h (181 ng/g versus 90.5 ng/g) and 24 h (171 ng/g versus 76.3 ng/g) **(Figure 5A)**. This pilot study suggests that LNPs enable JNJ64619178 to reach brain concentrations close to the IC50 identified in our *in vitro* studies. Thus, a novel LNP formulation further improved JNJ64619178 brain accumulation, indicating promising potential to cross BBB and target brain tumors

**Figure 5.**
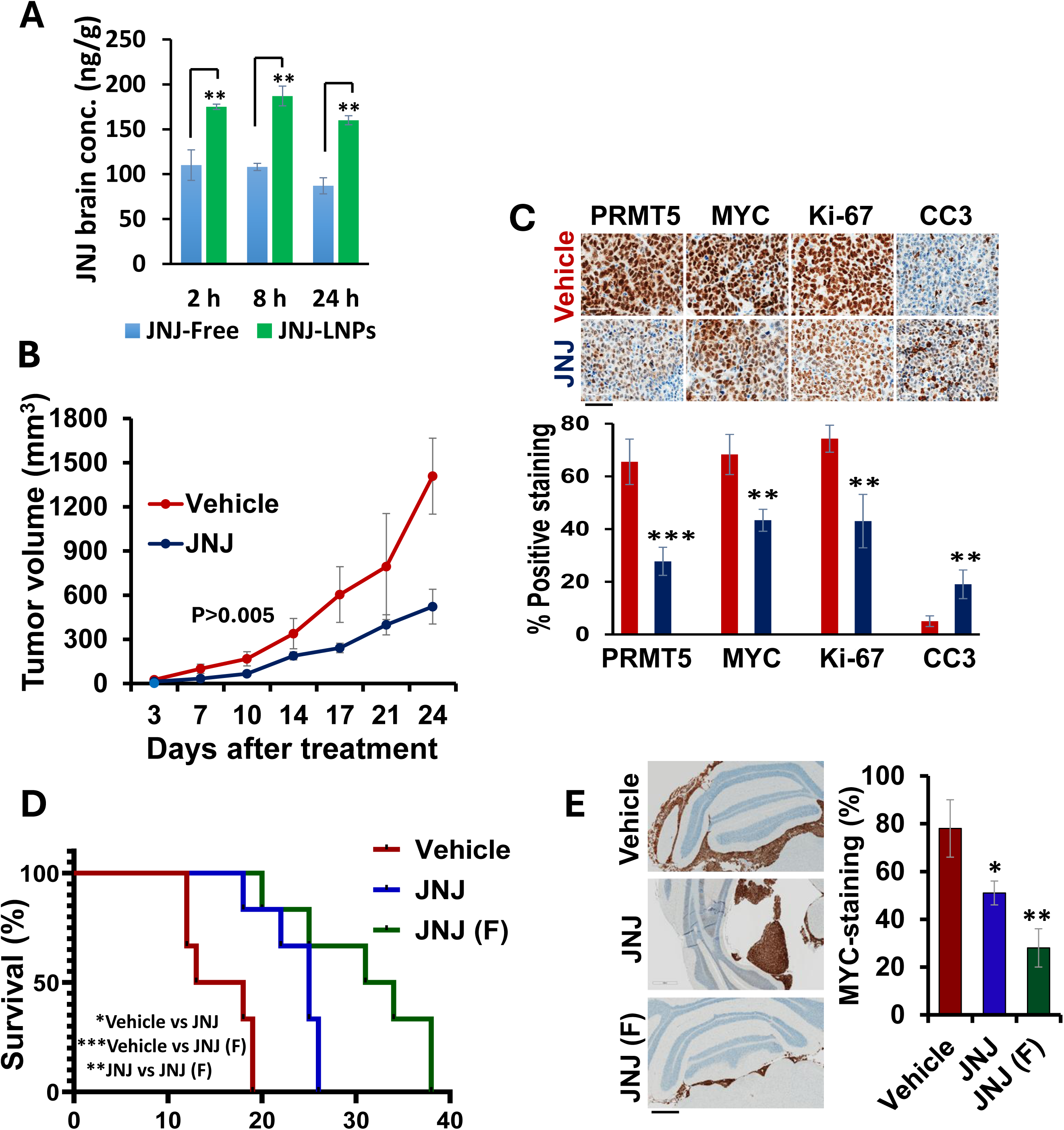
*In vivo* effects of JNJ64619178 (JNJ) in MYC-amplified MB-bearing mice. **(A**) JNJ brain concentration in BALB/c mice (N=3) following oral administration of 10 mg/kg JNJ at different timepoints. **p<0.01 (Student’s t-test). **(B)** NSG mice (N=5) with subcutaneously xenografted HD-MB03 cells were treated orally with vehicle or JNJ (10 mg/kg) five times a week for three weeks. Tumor volume measurement of xenografted mice following treatments. The differences noted between treatment groups show comparison by Student ‘s t-test of the tumor volumes on 21 days post treatment (p<0.005). **(C)** Representative IHC images (40 × magnification with 60 µm scale bar) of PRMT5, MYC, Ki-67, and CC3 expression in xenografts 21 days post-treatment, as indicated. Bar graphs below show the percentages of PRMT5, MYC, Ki-67 and CC3 positive cells derived from immunohistology scores, which were semi-quantitated in the tumors of three xenografted mice. **p<0.01; ***p<0.005 (Student’s t-test). **(D)** NSG mice (N=6) with orthotopically xenografted HD-MB03 cells were treated daily with vehicle or JNJ (10 mg/kg) or JNJ (F) (10 mg/kg) for two weeks. Survival analysis of xenografted mice using Kaplan-Meier (long-rank test). *p<0.05, **p<0.01, ***p<0.001. **(E)** Representative IHC images (4x magnification with 600 µm scale bar) and respective quantification showing MYC-positive tumors in the mouse cerebellum. The percentage of MYC, derived from immunohistology scores, was semi-quantitated in the tumors of three xenografted mice 21 days post-treatment. *p< **p<0.01 (Student’s t-test).

To examine the anti-tumor efficacy of JNJ64619178, we first employed a subcutaneous mouse model of MYC-driven MB where NSG mice were subcutaneously transplanted with MYC-amplified (HD-MB03) MB cells and treated with vehicle control or JNJ64619178. JNJ64619178 treatment led to an approximately 60% inhibition of tumor growth relative to vehicle control (**Figure 5B**), underscoring its therapeutic potential against MYC-driven medulloblastoma. In addition, treatment with JNJ64619178 did not cause significant differences in body weight between control and treatment groups **(Supplementary Figure 4A)**, suggesting the tolerability of JNJ64619178 in mice. We next determined the effect of JNJ64619178 on the expression of PRMT5, MYC and Ki-67 (proliferation marker) in xenografted tumors. Immunohistochemical analysis showed that JNJ64619178 significantly reduced the expression of PRMT5, MYC and Ki-67 **(Figure 5C)**. These data suggest that JNJ64619178 not only reduced tumor growth but also inhibited target proteins PRMT5 and MYC and cell proliferation in subcutaneous xenografted tumors.

To extend our findings from subcutaneous models, we investigated the antitumor efficacy of JNJ64619178 in orthotopic xenografts generated via intracerebellar injection of HD-MB03 cells into NSG mice. Ten days post tumor injection, mice (*n* = 6 per group) were treated with vehicle or JNJ64619178 free or JNJ64619178 formulated with LNPs. Treatment with JNJ64619178, either free or formulated as a lipid nanoparticle (LNP), significantly extended the survival of mice, with the formulated version showing efficacy, compared to vehicle control (**Figure 5D)**. These results were consistent with reduced tumor growth and MYC expression in the cerebellum **(Figure 5E)**. This finding suggests that when LNP-formulated, JNJ64619178 has improved brain-penetrance and thereby greater anti-tumor efficacy in MB. We observed no obvious toxicity of LNP-formulated JNJ64619178 in mice, as there were no significant changes in the histopathology of vital organs and body weight between free and formulated treatment groups **(Supplementary Figure 4B and C)**. Together, *in vivo* results suggest the anti-MB potential of JNJ64619178 not only against subcutaneous tumors but also orthotopic tumors.

## DISCUSSION

The minimal improvement in survival of Group 3 MB patients who receive current therapies points to a need for novel targeted therapeutic approaches ^6^. MYC amplification or overexpression is a critical determinant for MB development, progression and therapy-resistance ^45^. Several studies have shown that MYC transcription and its associated oncogenic programs play a key role in these events ^46^. However, how MYC transcription itself is regulated, particularly at the epigenetic level, remains poorly understood. We previously reported that PRMT5, an epigenetic enzyme, stabilizes MYC oncoprotein by physically interacting with it in MYC-driven (Group 3) MB ^28^, but it is unexplored if PRMT5 can regulate MYC transcription epigenetically as well. Here, we provide evidence showing that PRMT5 is a novel epigenetic activator of MYC transcription in MB. First, knockdown of PRMT5 specifically inhibited MYC mRNA and its subsequent transcriptional activity. Second, PRMT5 binds to the proximal promoter region of the MYC gene along with BRD4 (BET protein). Third, H4R3me2s is highly enriched on the proximal promoter region of the MYC gene. Fourth, PRMT5 inhibition suppresses MYC-induced transcriptional programs and alternate splicing events associated with energy metabolism. Fifth, pharmacological inhibition of PRMT5 by small molecule inhibitors specifically inhibited MB cell growth in MYC-dependent manner. Finally, pharmacological inhibition of PRMT5 strongly reduced the tumor growth in MB xenografted mice by downregulating MYC expression. Together with our previous findings, our data highlight that PRMT5 not only regulates MYC expression at the post-translation level, but also at the transcription level.

Gene expression is a tightly regulated process that involves the participation of multiple transcriptional regulatory proteins such as transcription factors, co-activators and co-repressors as well as chromatin-modifying enzymes. Consistent with the fact that BET protein family member BRD4 is the major epigenetic regulator that activates MYC transcription in cancer cells ^35^, we indeed confirmed that BRD4 binds to the MYC promoter and regulates MYC transcription in MYC-amplified MB cells. Interestingly, PRMT5 interacts with BRD4 that is bound to the MYC promoter, suggesting that BRD4 may recruit PRMT5 to the MYC promoter. Our findings also suggest that PRMT5-mediated symmetric dimethylation of H4R3 could also activate the transcription of MYC when BRD4 is recruited to the MYC promoter. Future studies focused on defining the epigenetic mechanisms governing MYC regulation may offer novel therapeutic avenues for targeting MYC expression.

PRMT5 has been shown to play a critical role in tumor maintenance by controlling gene expression, the epigenetic landscape, mRNA processing/splicing and oncogenic pathways ^47^. We demonstrated that PRMT5 inhibition in MYC-amplified MB cells leads to significant remodeling of the MYC transcriptome and also significantly impairs spliceosome function, leading to broad disruption of alternative splicing events (ASEs), particularly in metabolism-associated pathways. It is worthwhile noting that such alterations, specifically including changes in ASEs, were also found in other cancer types following PRMT5 inhibition ^37,38^ indicating that these molecular changes are bona fide consequences of PRMT5 inhibition. Dysregulated pre-mRNA splicing and metabolic reprogramming constitute fundamental hallmarks of MYC-driven cancers. Both PRMT5 and MYC have been shown to regulate pre-mRNA splicing, leading to metabolic alterations in cancer cells ^48^. Collectively, our data identify PRMT5 as a central hub that functionally links aberrant splicing and metabolic reprogramming in MYC-driven medulloblastoma. This study identified a downstream mechanism of PRMT5 that reprograms metabolism in MYC-driven MB cells by regulating mRNA splicing.

Despite promising preclinical activity, many new therapeutic agents for medulloblastoma fail in clinical trials owing to insufficient BBB penetration. Previous work demonstrated that LLY-283, a potent PRMT5 inhibitor, efficiently crosses the BBB and prolongs survival in orthotopic glioblastoma xenografts ^37^. Here, we show that JNJ64619178, a more potent and clinically relevant PRMT5 inhibitor (ClinicalTrials.gov NCT03573310), also crosses the BBB and exhibits enhanced antitumor activity against orthotopic MYC-driven MB, particularly when delivered via lipid nanoparticle formulation. To our knowledge, this is the first demonstration of superior antitumor activity of a nanoformulated PRMT5 inhibitor relative to its free molecule formulation in a brain cancer setting. In addition, JNJ64619178 significantly prolonged survival in mice with orthotopic Group 3 MYC-driven medulloblastoma, highlighting its therapeutic promise. Nevertheless, further preclinical evaluation of clinically relevant PRMT5 inhibitors is warranted.

Our study provides important mechanistic insight linking PRMT5 inhibition to the suppression of MYC-driven transcriptional programs, the disruption of constitutive splicing, impairing proliferation and survival in MB and providing a strong rationale for targeting PRMT5 in MYC-driven MB. Our proof-of-concept in vivo data using a brain-penetrant chemical probe support the feasibility of brain-penetrant PRMT5 inhibition as a therapeutic strategy. Based on our findings on the impact of PRMT5 inhibition on metabolic activities, assessing the efficacy of PRMT5 inhibitors in combination with metabolic inhibitors is warranted. Moreover, considering that Group 3 MB patients exhibit minimal therapeutic benefit from chemoradiation treatment and studies in other cancers have identified PRMT5 inhibition as a chemo- or radio-sensitization target [49], combining PRMT5 inhibition with existing chemotherapy (e.g., cisplatin) and radiotherapy regimens may enhance therapeutic outcomes for this patient population. ^49^Together, our findings highlight the importance of further evaluating PRMT5 inhibition in patient-derived orthotopic xenograft models as a critical step toward clinical translation.

## Ethics Statement

All animal experiments were performed in compliance with protocols approved by the University of Nebraska Medical Center Institutional Animal Care and Use Committee (IACUC).

## Authorship Statement

NKC conceptualized and designed the study. DK, NKC, AS, AKD, RK, LD, YSC, SS, GN, and SR performed the experiments and analyzed the data. NKC, DK, and AKD wrote the manuscript. BG, HB, DJM, and DWC reviewed the manuscript and contributed significantly to the interpretation of the data. All authors read and approved the manuscript.

## Data Availability Statement

All data that supports the findings of our study are available from the corresponding author upon request.

## Acknowledgements

The authors thank the Child Health Research Institute and the Fred and Pamela Buffett Cancer Center supported (Genomics, Flow Cytometry, Bioimaging and Tissue Sciences) core facilities at UNMC. The authors thank Matthew Sandbulte, PhD, of the Child Health Research Institute at Children’s Nebraska and the University of Nebraska Medical Center for his help in editing this manuscript. Graphical abstract was created using Biorender.

