## Supplementary Figures 1-4 for "PRMT5 as an Epigenetic Target for Group 3 (MYC-driven) Medulloblastoma"

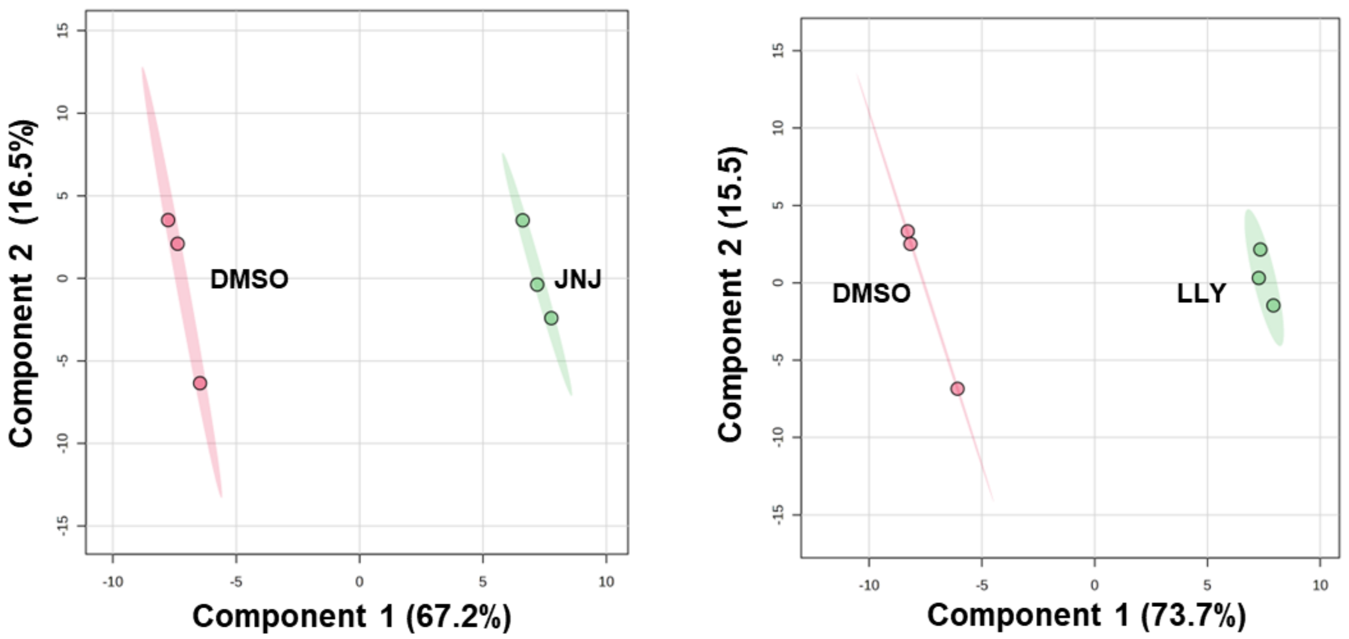


**Supplementary Figure 1. Principal Component Analysis (PCA) between DMSO and JNJ64619178 or LLY-283 treatment groups.** PCA plot showing the segregation of vehicle (DMSO) and JNJ64619178 (1 µM for 24 h) or LLY-283 (5 µM for 24 h) treated HD-MB03 cells based on their metabolite profiles. Each colored circle represents a biological replicate of the treatment condition. Component 1 indicates the degree of variation between the groups based on their total metabolite content, and component 2 indicates the differences within the groups.

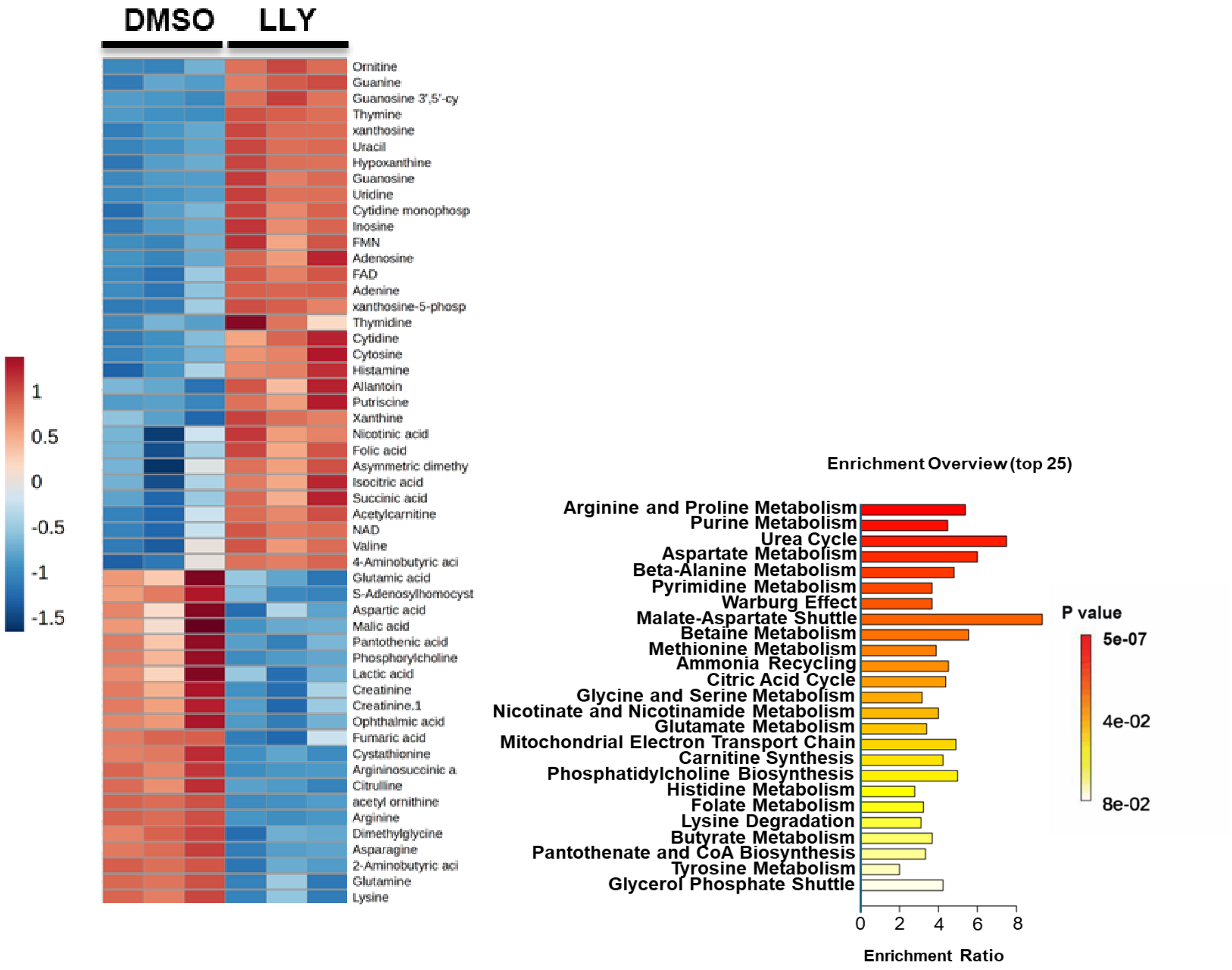


**Supplementary Figure 2. LLY-283 (LLY) treatment alters energy metabolism in MB. (A)** Heatmap showing top 60 metabolites altered in HD-MB03 cells treated with LLY (5 µM) in triplicate for 24 h. Color intensity represents the magnitude of alteration in individual metabolites. Scale for color intensity is shown adjacent to the heatmap. **(B)** Pathway analysis showing significantly altered metabolic pathways in LLY treated HD-MB03 cells, compared to DMSO solvent. Scale under the pathway plot shows the fold enrichment and color scale adjacent to pathway plot indicates significance (*p*-value) for altered pathways.

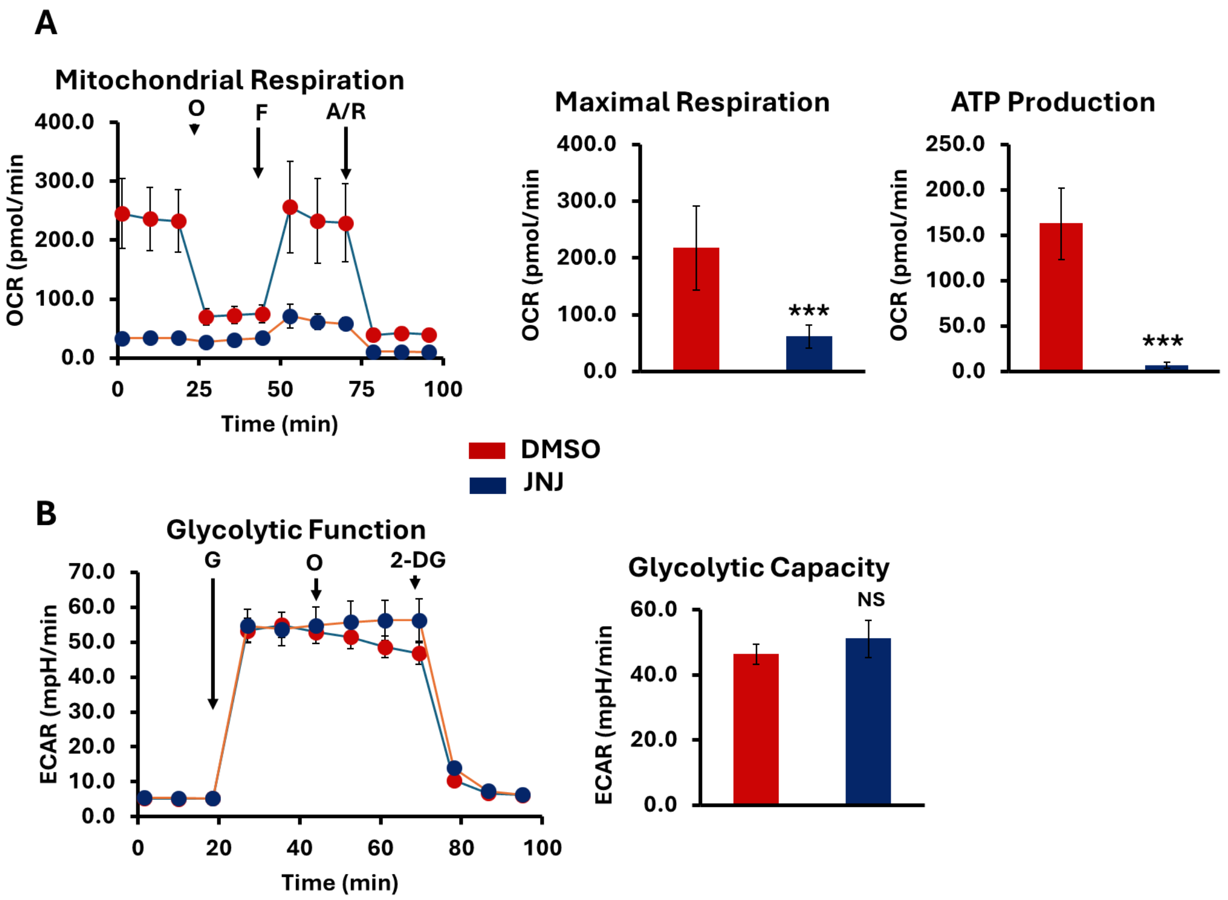


**Supplementary Figure 3. JNJ64619178 (JNJ) treatment alters energy metabolism in MB. (A)** Oxygen consumption rate (OCR) analysis for mitochondrial oxidative phosphorylation status in D-425 cells after treatment with 1 μM JNJ for 24 h. O, oligomycin; F, FCCP; A/R, antimycin/rotenone. The bar graphs show maximal respiration and ATP production activities derived from OCR activities shown in line graph. The results represent the mean ± SEM of three replicates. ***$p < 0.001$ (Student t test, DMSO vs JNJ). **(B)** Extracellular acidification rate (ECAR) analysis for glycolytic activities in D-425 cells after treatment with 1 μM JNJ for 24 h. G, glucose; O, oligomycin; 2-DG, 2-deoxyglucose. The bar graph shows glycolytic capacity derived from ECAR activities shown in line graph. The results represent the mean ± SEM of three replicates. NS (not significant).

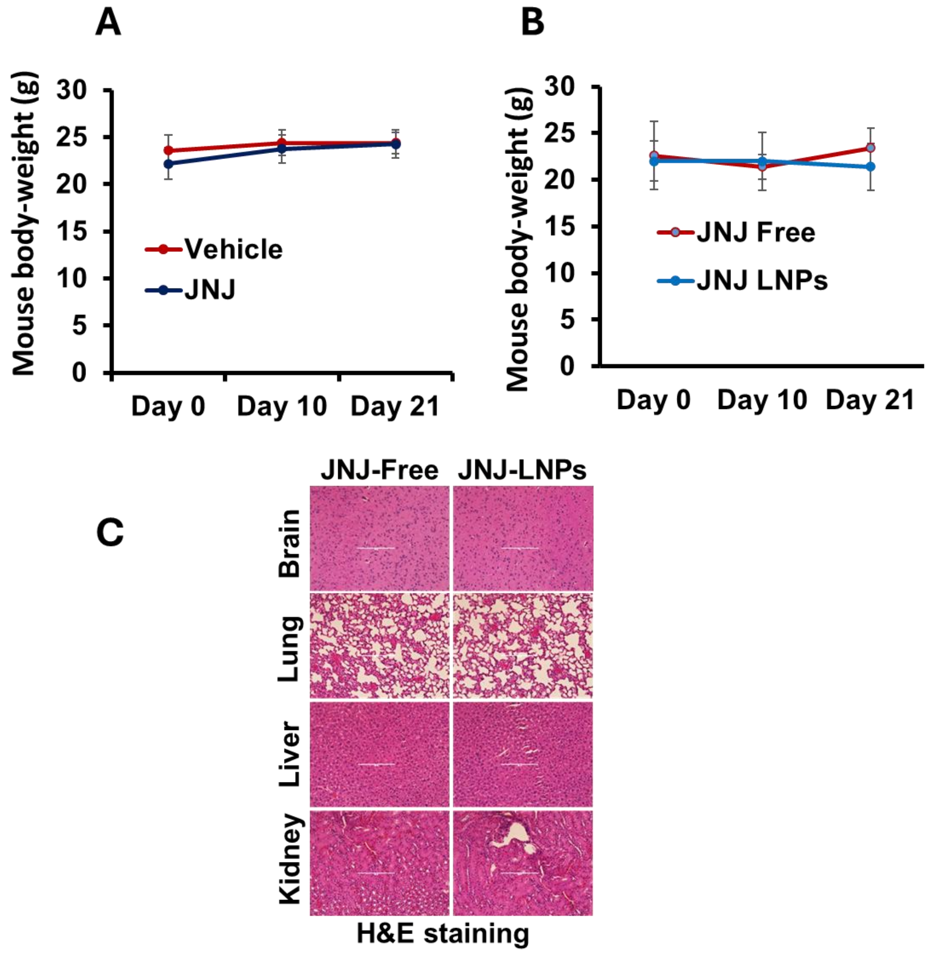


**Supplementary Figure 4. Effects of JNJ64619178 (JNJ) on body weight and histology of the MB xenograft mice. (A)** The mean body weight of mice following treatment with vehicle or JNJ as indicated. **(B)** The mean body weight of mice following treatment with JNJ-free or JNJ-LNPs (formulated) as indicated. **(C)** Histopathology (H&E) of the vital organs of MB xenografts following 21 days post treatment with JNJ-free or JNJ-LNPs (formulated). The images were scanned and captured using digital scanner EVOS Image system at 20x magnification.
